# Optimization for Predictive Gene Circuit Design Automation

**DOI:** 10.1101/2022.06.21.497000

**Authors:** Frederick Starkey, Filippo Menolascina

## Abstract

Synthetic Biology aims to rationally engineer biological systems. Current methods often employ an initial human designed circuit topology and utilise iterative approaches, e.g. directed evolution, to fine-tune part function. This approach can be extremely time consuming and resource intensive whilst often reaching sub-optimal solutions. A design workflow in which circuits and parts are designed in silico can overcome such limitations. Here we describe a method to automatically design synthetic gene circuits with user-specified dynamics; unlike some previous contributions our algorithm is able to design circuits with analog, not just digital behaviours. We demonstrate the capabilities of our approach benchmarking it on a number of different gene circuits design tasks. We review and compare the performance of our method against state of the art and outline future opportunities for development. Finally, to foster adoption, we make our algorithm available through a web app.

## Background

The design of gene circuits aims to produce biological organisms which can be controlled by arbitrarily complex programmable logical functions. At present, workflows for the design and construction of gene circuits require many iterations even for the most simple circuits. In cases where rational design methodologies are applied (Elowitz et al. [1]) physical models produced are specific to the function of the device in question. As our ability to build larger, more complex gene circuits expands, for example thanks to the availability of Biofoundries [27], scalable circuit design techniques become the bottleneck in the process of advancing synthetic biology towards applications.

The evolution of electronic design automation (EDA) has been seen as painting a potential roadmap for progress in gene circuit design automation. Prior to the adoption of EDA in the 1970s, the design of integrated electronic circuits was a laborious process. Circuits were constructed manually based on hand-drawn designs. The advent of EDA and creation of hardware description languages such as VHDL and Verilog to describe the topology and behaviour of electronic circuits catalysed the widespread production of complex electronic circuits. In turn, this paved the way for the modern information age. A number of authors in Synthetic Biology have now advocated for BioDesign Automation (BDA) or Gene Circuit Design Automation (GCDA), a biological equivalent of EDA aimed at facilitating a similar leap forward in genetic engineering realising the promise of Synthetic Biology.

Figure 1 demonstrates that whilst it is tempting to aim to apply EDA techniques to gene circuit design automation in a 1-to-1 manner, there are important functional differences between electronic and genetic circuits which hinder such an endeavor.

**Figure 1:**
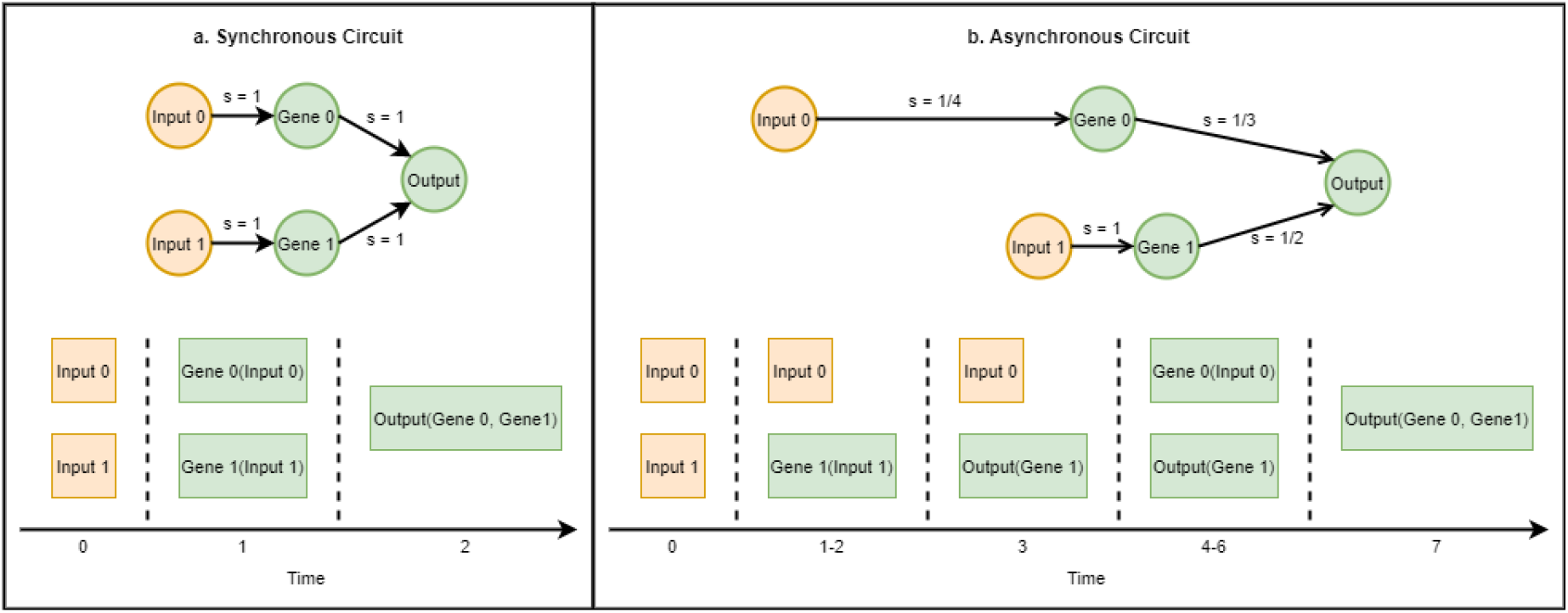
Synchronous and Asynchronous Circuits. a. In synchronous circuits state changes of components are synchronised by a regular clock signal. Hence the speed of signal transmission (s) and time for a signal to pass from one component to another is uniform. b. Gene Circuits are asynchronous circuits in which the speed of signal transmission (s) is determined by chemical interactions and hence highly irregular. It is very unlikely that signals pass between components at the same time.

**Figure 2:**
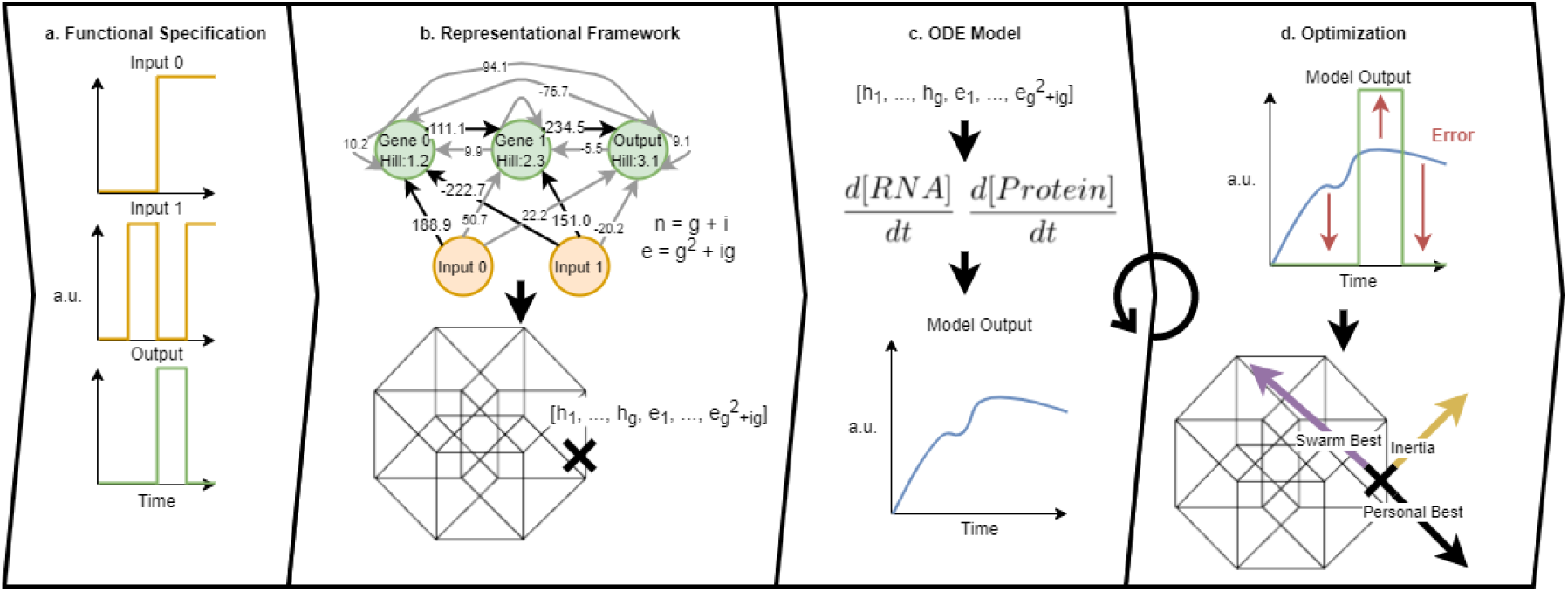
Typical workflow of our approach. a. The target functionality of the gene circuit is specified by providing time series for the input and output signals. b. A weighted network graph represents the regulatory interactions of a potential solution. Edge weights greater than 100 indicate an activation, less than -100 indicates a repression otherwise no interaction is present. Gene nodes each have a hill coefficient indicating the extent of cooperative binding in their regulation. Such a graph can be encoded as a position in *g* + *g*^2^ + *ig* dimensional space where *g* is the number of genes and *i* the number of input signals. c. A position in *g* + *g*^2^ + *ig* dimensional space is used to parameterize a system of ordinary differential equations which models the dynamic behaviour of a gene circuit. d. Particle Swarm Optimization is used to explore new positions and hence new gene circuits. Based on the difference between circuit behaviour and target output particles move through the search space according to the best known position across all particles, an inertia towards the particles previous position and the best position an individual particle has previously inhabited. Steps c. and d. are applied for each particle in each iteration of the optimization before the gene circuit encoded by the best particle position explored is returned as a solution.

**Figure 3:**
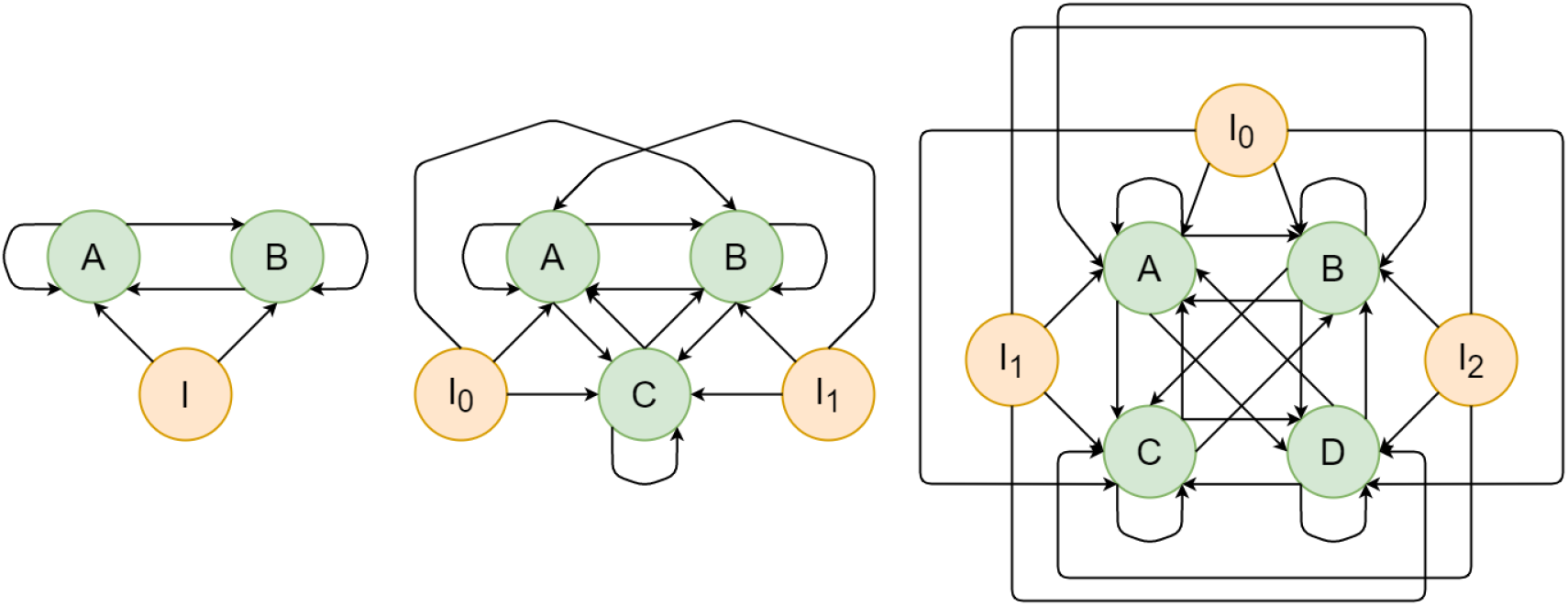
Regulatory Interactions as Network Graphs. Example topologies for network graphs encoding the regulatory interactions in gene circuits. Candidate gene circuits are parametizations of graphs of this structure containing *G* + *I* nodes and *G*^2^ + *IG* edges.

**Figure 4:**
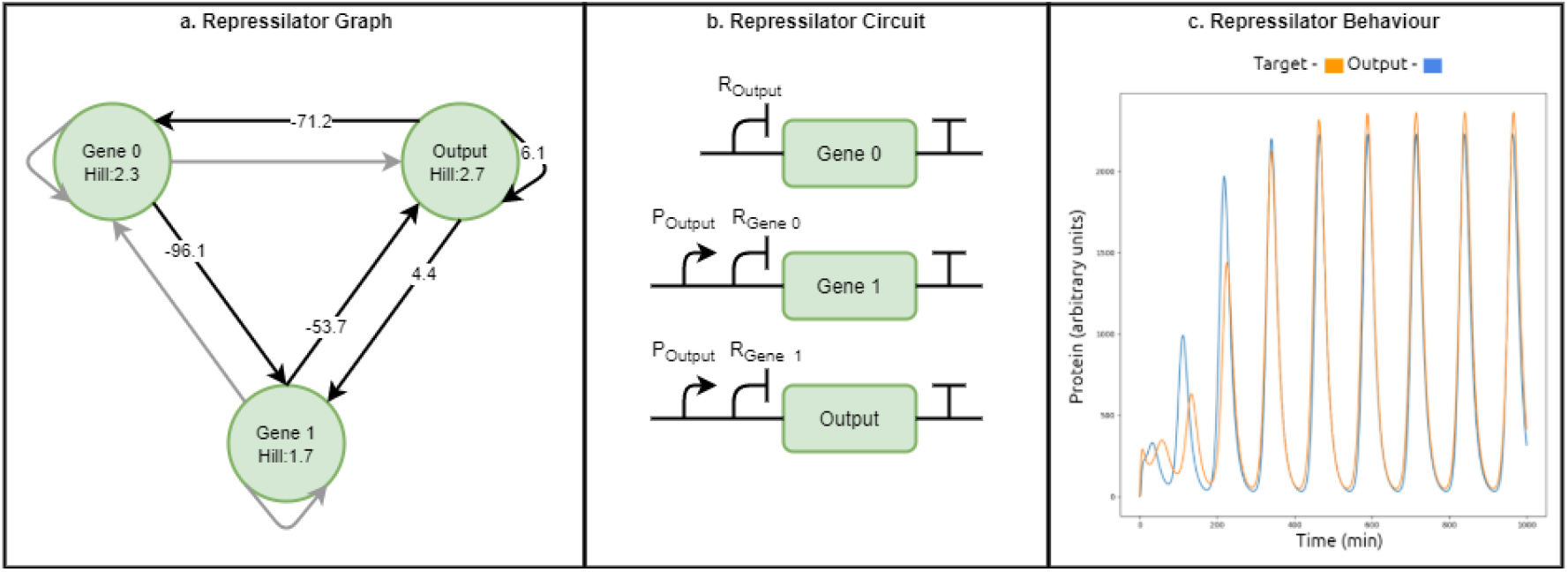
Repressilator. a. Network Graph representation of the Repressilator design. Green nodes represent genes each with a Hill coefficient. Edges represent potential regulatory interactions. Grey edges are unused in the circuit design. Black edges are parametized by a Michaelis Mentens constant - negative for repression and positive for activation. b. Parts scale circuit representation of Repressilator. The circuit consists of 3 genes with expression controlled by activators and repressors as outlined in a. c. Behaviour of the Repressilator design vs the target from Elowitz et al. [1]. Time series in blue is the output of the ODE model for the generated circuit design. Time series in orange is the output of the ODE model from Elowitz et al. [1]

**Figure 5:**
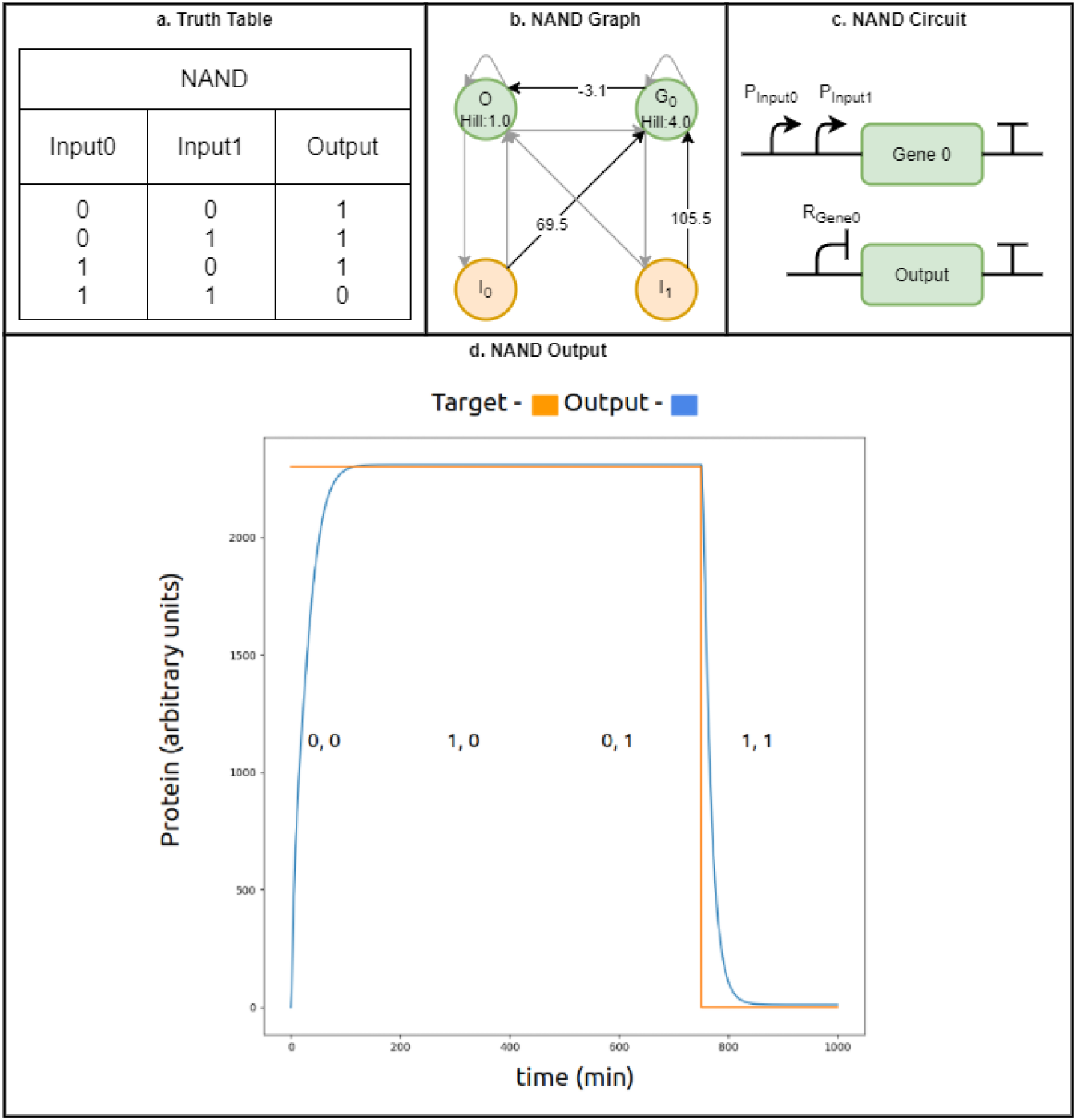
NAND Gate. a. Truth Table for NAND operation. Outlines the expected logical output for each possible pair of logical inputs.a. Network Graph representation of the NAND Gate design. Green nodes represent genes each with a Hill coefficient. Edges represent potential regulatory interactions. Grey edges are unused in the circuit design. Black edges are parametized by a Michaelis Mentens constant - negative for repression and positive for activation.c. Parts scale circuit representation of NAND Gate. The circuit consists of 2 genes with expression controlled by activators and repressors as outlined in a.d. Behaviour of the NAND design vs the target. Time series in blue is the output of the ODE model for the generated circuit design. Time series in orange is the target output for each pair of inputs in sequence as denoted by labels.

**Figure 6:**
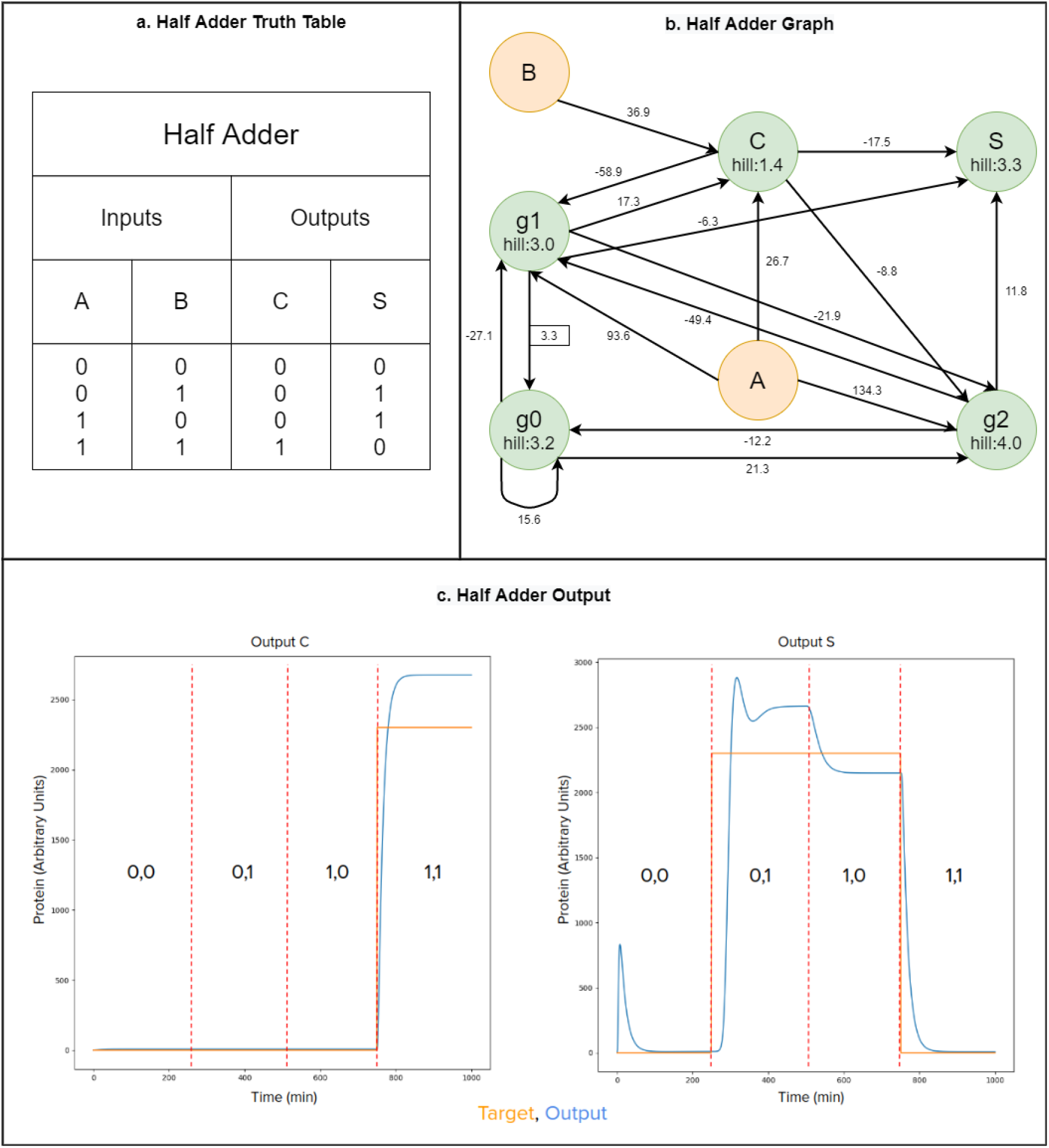
Half Adder. a. Truth Table for Half Adder. Outlines the expected logical output for each possible pair of logical inputs.a. Network Graph representation of the Half Adder design. Green nodes represent genes each with a Hill coefficient. Edges represent potential regulatory interactions. Edges are parametized by a Michaelis Mentens constant - negative for repression and positive for activation. c. Behaviour of the Half Adder design vs the target. Time series in blue is the output for genes C and S of the ODE model for the generated circuit design. Time series in orange is the target output for genes C and S for each pair of inputs in sequence as denoted by labels.

Electronic circuits are Synchronous Circuits in which state changes of components are kept in time by a regular clock signal. This paradigm has the benefit that individual components are context independent. For example in the electronic context an XOR Gate and an AND Gate taking the same input signals would provide their output at the same time. By contrast gene circuits are Asynchronous circuits, state changes of components (genes products) are determined by the kinetics of upstream biochemical/regulatory interactions that, in general, take place at different speeds, therefore making the “synchronous” assumption is inappropriate in GCDA. Considering again the XOR and AND gate example in the biological context; logically minimal gates based in a synchronous paradigm are not equivalent to functional gates in an asynchronous paradigm. Functional logic gates in the gene circuit context are likely to have a counter-intuitive topology which accounts for mismatches in signal transmission between parts.

Gene Circuit Design Automation became an active area of research over the past decade. Previous works in this field adopted a variety of different approaches to automatically design biological networks with user defined dynamic behaviour. Francois et al. [2] used an evolutionary Genetic Algorithm-like approach to design both bistable switches and oscillators. Otero-Muras et al. [3] formulated GCDA as a Mixed Integer Nonlinear Programming problem and provided a Matlab-based toolkit which they use to design a number of gene circuits based on a built-in parts library. Cello [4] seeks to borrow from EDA and allows users to define gene circuits in Verilog code within a web app. The Cello system is demonstrated by implementing 60 designed circuits in Escherichia coli with a 92% success rate when built in the wet lab. Our work draws inspiration from Hiscock et al. [5] who developed Genenet a gradient descent-based Python module with an emphasis on speed which utilises Theano [6] and Tensorflow [7] to optimize a user defined ODE system. In comparison to the work of Francois et al [2] we demonstrate that Particle Swarm Optimization converges to a solution more quickly than a Genetic Algorithm in Gene Circuit Design Problems. Whilst Otero-Muras et al. [3] and Cello [4] simplify the design problem by working from a built-in parts library even the largest libraries contain parts covering only a negligible section of the possible sequence space and hence limit the types of circuits which can be generated. Finally, the methods used by Otero-Muras et al. [3], Cello [4] and Hiscock et al. [5] all require complex computer programming knowledge which render them inaccessible to non-technical users.

Motivated by a comparison between the performance of different optimization techniques we build on the body of previous work using Global Particle Swarm Optimization (PSO) [8] to optimize gene circuits *in silico* for a user defined function. To allow access to the full sequence space of parts, our algorithm provides parameters describing the behaviour of parts perfectly calibrated to produce a specific circuit rather than working from a pre-characterised library. Crucially, the optimization process we propose is able to identify rather counter intuitive solutions which circumvent the signal mismatching problem inherent to asynchronous genetic circuits. We demonstrate that PSO outperforms other techniques on this optimization problem. Whilst previous work in GCDA mainly catered to a computer science audience, here we aim to increase adoption, making our program available via a graphical user interface packaged in a web app. In the following we present our solution, along with some example circuit designs.

## Methodology

### Modelling Gene Circuits using Ordinary Differential Equations

As a modelling formalism for the circuits our method will design, we selected Ordinary Differential Equations (ODE) which can be adapted to model a circuit with arbitrary regulatory interactions. Although it is computationally intensive to evaluate the results of many ODE systems we chose to pursue this methodology given the highly descriptive nature of the output of the candidate circuits in both static and dynamic phases. Transcription and translation for each gene in a candidate solution are formalised as 2 ODEs - one for production of RNA and one for production of protein.

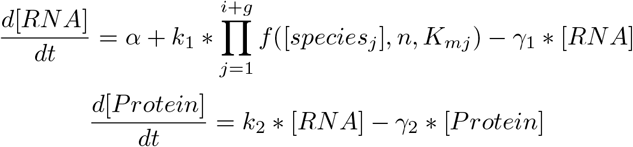

*α* is used to model promoter leakiness and gives the transcription rate irrespective of regulation. *k*_1_ and *k*_2_ account for the basal transcription and translation rate respectively. Similarly *γ*_1_ and *γ*_2_ represent the RNA and protein degradation rates.

Function *f* uses Michaelis Mentens kinetic to model the regulatory effect of each input species (provided as a time series by the user) and of *g* genes in the candidate gene circuit. Function *f* is formalised as a piecewise version of the Hill function which compactly represents both activation and repression.

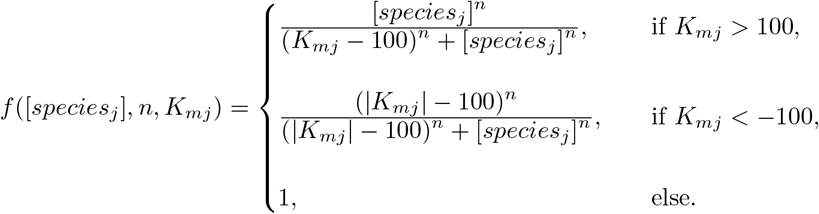

[*species*_*j*_] gives the quantity (in arbitrary units) of some regulator (either input chemical or protein) at single time point. *n* gives the Hill coefficient describing the degree of cooperative binding of the ligand [*species*_*j*_]. For our purposes we bound *n* between 1 and 5. These bounds ensure a state of negative cooperative binding in which affinity for ligand binding decreases as more molecules bind to the promoter. *K*_*mj*_ encodes both the type of regulatory interaction (activation if greater than 100, repression if less than -100 and else no interaction) and the Michaelis Mentens constant for the regulation. *K*_*mj*_ is bounded between -300 and 300. When greater than 100 *K*_*mj*_ − 100 represents the concentration of activator to express at half the maximal rate, similarly when less than -100 |*K*_*m*_*j*| *−* 100 represents the concentration of repressor to express at half the maximal rate. In the range -100 to 100 inclusive *K*_*m*_*j* encodes that no regulatory interaction exists.

The design of a gene circuit for a user defined function is hence the search for such an ODE system which defines behaviour matching as closely as possible the target output signal.

### Regulatory Interactions as a Network Graph

The set of regulatory interactions which characterise a candidate gene circuit can be encoded as a network graph. A gene circuit with *G* genes and *I* input signals will contain *G* + *I* nodes, each of the *G* gene nodes will have a parameter encoding its Hill coefficient. To represent all the regulatory interactions possible between *G* genes and *I* input signals the gene circuit graph must contain *G*^2^ + *IG* edges. Each edge from node *A* to node *B* represents the regulation of gene *B* by species *A* (either input signal or gene). Edges in the gene circuit graph are weighted by *K*_*mj*_ which encodes the kind of regulatory interaction and that interactions Michaelis Menten constant.

Given this formalisation for representing the regulatory interactions of arbitrary candidate gene circuits, any gene circuit can be encoded as a position in *G* + *G*^2^ + *IG* dimensional space. The search for an ODE system which defines gene circuit behaviour matching as closely as possible the target output signal can hence be defined as a search through this *G* + *G*^2^ + *IG* dimensional space.

### Optimization Methods

We adopt an optimization to minimize an error function which accounts for the distance between the output signal of a candidate gene circuit and (as evaluated by solving a system of ODEs) the number of regulatory interactions in a solution.

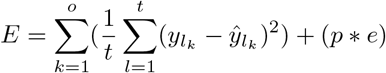

The error (*E*) is the sum of the average square distance over each time point for each output signal plus a regularisation factor for each regulatory interaction (to encourage minimal solutions).

After a performance comparison between a number of potential approaches (Supplementary) we chose to use Particle Swarm Optimization (PSO) to solve the gene circuit design problem. We begin by defining a swarm of particles each with a position in *G* + *G*^2^ + *IG* dimensional space encoding a candidate gene circuit. The ODE system for each particle (solution) is then solved: the error function for each solution is evaluated at this stage. Based upon the evaluation of the candidate solutions in the swarm the position (Eq. 2.) and velocity (Eq.1.) are iteratively updated to generate new and improved candidate gene circuits.

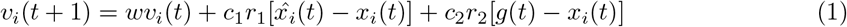

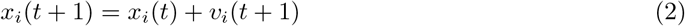

The position *x*_*i*_ of particle *i* is updated in accordance with the velocity *v*_*i*_. The velocity *v*_*i*_ of particle *i* is updated according to the sum of *wv*_*i*_(*t*) - an inertia towards the particles previous velocity, 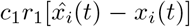 - a randomly weighted cognitive factor towards the best position ever inhabited by particle *i* and *c*_2_*r*_2_[*g*(*t*) − *x*_*i*_(*t*)] - a randomly weighted social factor towards the best position ever inhabited by any particle in the swarm.

## 1 Results

We detail here some example circuit designs for a variety of functions. The repressilator is a simple circuit with an oscillating output, it is frequently used as an example of how modelling can be integrated into the gene circuit design process. The NAND gate is a circuit with two inputs that performs a simple logical operation. The NAND operation is of particular interest due to being functionally complete (i.e. any logic function can be implemented used only NAND gates); additionally it allows us to demonstrate our techniques ability to reach a minimal solution to a design problem. To demonstrate scalability we also design a half adder circuit which outputs the sum of its two inputs.

### Repressilator

The repressilator proposed by Elowitz et al.[1] demonstrates how an oscillating output can be obtained using a minimal circuit of 3 genes. To test the ability of our approach to reconstruct the repressilator we implemented the ODE model as described by Elowitz et al [1] and used the output signal as the target for the optimization process as described in methods.

The repressilator circuit was used to fine-tune the hyperparameters of our optimization. Inertia *w* was tested between 0.1 and 1 sampling every 0.1. The best inertia setting found was 0.3. Cognitive and social parameters *c*_1_ and *c*_2_ were tested as a pair based on the heuristic that they should always sum to 4. We varied each of *c*_1_ and *c*_2_ between 1 and 3 sampling every 0.1. The best cognitive factor setting was found to be 1.3 and social factor setting 2.7. The penalty factor for edges *e* was sampled every 250 varying between 2000 and 3000. The best penalty value was 2750. To prevent repetition of the laborious process of fine-tuning hyperparameters these settings producing the lowest error on the repressilator were used for all subsequent circuit designs.

The criteria for a successful repressilator circuit design was to produce oscillations with approximately the same magnitude and frequency as the target. We were unable to achieve this behaviour with any less than 75 particles and 500 iterations.

### NAND Gate

Unlike the repressilator (analog signal) this biological equivalent of a fundamental electronic component is digital in nature: this fundamental difference with Elowitz’ circuit made us choose the automatic design of a NAND gate as a complementary demonstration of the capabilities and flexibility of our approach.

The successful NAND gate design uses fine-tuned hyperparameters from the repressilator. Inertia 0.3, Cognitive factor 1.3, Social factor 2.7 and Edge Penalty 2750.

There are a number of criteria for a successful NAND gate design. Upon receiving a new input signal the circuit must quickly assume the appropriate output (ideally within 100 minutes - half the size of each input window). When the circuit is in the off state there must be close to 0 protein expression - bearing in mind a true 0 value is not possible due to leakiness of repression and activation. The output for each of the input signals corresponding to a on circuit state should ideally be the same. However as long as there is a significant difference between the on and off circuit states we do not particularly mind if the target output magnitude of 2400 in the on state is reached. We were unable to achieve a design with the target behaviour using any less than 75 particles and 500 iterations.

### Half Adder

As we are motivated to provide the synthetic biology community with a gene circuit design algorithm able to in silico synthesise complex gene circuits we sought to test its performance on a more complex problem: the design of a half-adder which outputs the sum of 2 bits.

The Half Adder design uses fine-tuned hyperparameters from the repressilator. Inertia 0.3, Cognitive factor 1.3, Social factor 2.7 and Edge Penalty 2750.

There are a number of criteria for a successful Half Adder design. Upon receiving a new input signal the circuit must quickly assume the appropriate output (ideally within 100 minutes - half the size of each input window). When the circuit is in the off state there must be close to 0 protein expression - bearing in mind a true 0 value is not possible due to leakiness of repression and activation. For the S output some settling time in the 0,0 input state is required for this criteria to be met. The output for each of the input signals corresponding to a on circuit state should ideally be the same. We were not able to recover this behaviour for the S output however given the clear differences between the on and off circuit states we still believe this constitutes a successful design. We were unable to achieve a design with acceptable behaviour using any less than 100 particles and 750 iterations. It is possible that a more optimal design could be found by further increasing the number of particles and iterations however the massive increase in complexity (due to the extra ODE system evaluations) required to do this made it infeasible given our computational resources.

## Conclusion

We present a system for the optimization of gene circuit designs. We show that this system can be used via a graphical user interface to design a variety of gene circuits requiring a time series for the input signals and expression of target species. By providing the user with parameters characterizing the dynamics of parts required for a specific circuit this Gene Circuit Design Automation strategy provides the first step in a bespoke predictive design workflow for Synthetic Biology. Using predictive design to generate gene circuit and parts specifications in silico facilitates a reduction in the number of wet lab iterations required to produce functional gene circuits. Such a streamlined predictive workflow for Synthetic Biology will allow the construction of larger circuits at lower cost in a reduced time. We provide our system in a user-friendly web app requiring no computer programming expertise in order to encourage adoption throughout the Synthetic Biology community. Although we demonstrate the performance of our system on gene circuits approaching the size limit of what is possible to construct with current wet lab techniques, future work could focus on improving the performance of our optimization techniques. In particular an examination of more complex swarm intelligence-based optimization algorithms is likely to yield promising results given the performance of the vanilla PSO algorithm. Nobile et al. [23] and Korenaga et al. [24] propose varients on the PSO algorithm designed to self tune hyperparameters and improve performance on high-dimensional problems respectively. The Bat Algorithm proposed by Xin-She Yang et al. [26] is another swarm based optimization technique which has been shown to outperform PSO on some benchmarks and is likely to perform well on the Gene Circuit Design Automation problem. By utilising our PSO based Gene Circuit Design Automation web app wet lab synthetic biologists generate and iterate on circuit designs with user defined dynamic behaviour. By integrating computational tools such as this into the gene circuit production workflow users can more rapidly produce functional gene circuits at a scale which would be infeasible for a non-computational approach.

